# Thermal Denaturation of DNA G-Quadruplexes and their Complexes with Ligands: Thermodynamic Analysis of the Multiple States Revealed by Mass Spectrometry

**DOI:** 10.1101/370254

**Authors:** Adrien Marchand, Frédéric Rosu, Renato Zenobi, Valérie Gabelica

## Abstract

As the idea that G-quadruplex nucleic acid structures are involved in cellular processes is gaining support, it becomes important to develop ligands that specifically target G-quadruplexes. However, ligand design is complicated because there are multiple G-quadruplex target sequences, some sequences are polymorphic, and very few ligand-quadruplex structures in solution were solved to date. Further, structure alone does not reveal the driving forces for ligand binding. To know *why* a ligand binds, the thermodynamics of binding must be characterized. Electrospray mass spectrometry makes it possible to detect and quantify each specific stoichiometry in terms of number of strands, number of specific cations, and number of ligands, and thus allows one to simultaneously determine the equilibrium constants for the formation of each complex. We designed and built a temperature-controlled nano-electrospray source to monitor thermal denaturation by mass spectrometry (“MS-melting”). We studied the thermal denaturation of G-quadruplexes, including the c-myc promoter and several telomeric sequence variants, and their complexes with popular ligands (Phen-DC3, TrisQ, TMPyP4, Cu-ttpy). From the temperature dependence of the equilibrium constants, we determined the enthalpic and entropic contributions to the formation of each stoichiometric state. In absence of ligand, we untangled the potassium-induced G-quadruplex folding thermodynamics, one potassium ion at a time. The formation of each quartet-K^+^-quartet units is strongly enthalpy driven, with entropy penalty. In contrast, the formation of quartet-K^+^-triplet units is entropically driven. For this reason, such misfolded structures can become more abundant as the temperature increases. In the presence of ligands, mass spectrometry also revealed new states at intermediate temperatures. For example, even in cases where only a 1:1 (ligand:quadruplex) is observed at room temperature, a 2:1 complex predominates at intermediate temperatures. Mass spectrometry also makes it easy to distinguish ligand bound to the 2-quartet structures (containing 1 K^+^), the 3-quartet structures (containing 2 K^+^) and to the unfolded strand (no specific K^+^). We confirm that TrisQ binds preferably, but not exclusively, to 3-quartet structures, Phen-DC3 binds to a 2-quartet structure, while the porphyrin ligand TMPyP4 is characterized as non-selective, because it binds to all forms including the unfolded one. The thermodynamics of ligand binding to each form, one ligand at a time, provides unprecedented detail on the interplay between ligand binding and changes in G-quadruplex topology.

**TOC Graphics:** 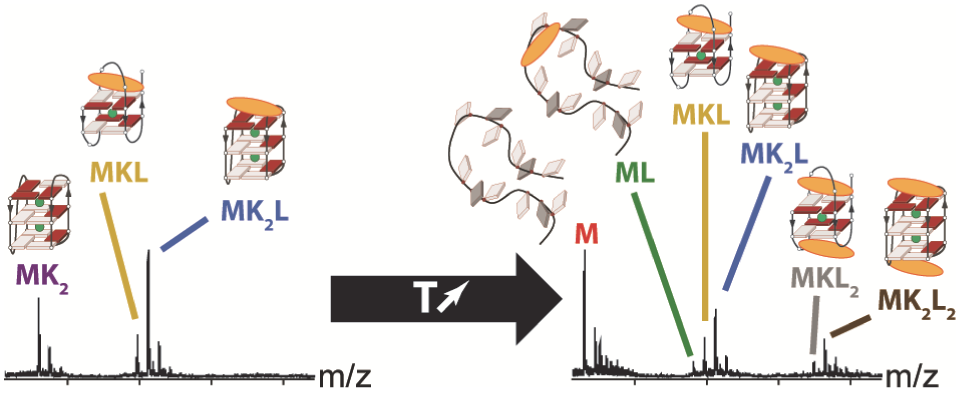

## INTRODUCTION

Besides storing genetic information, nucleic acids are also involved in the regulation of gene expression. To ensure such function, they adopt particular three-dimensional structures, one of which being the G-quadruplex:1 a four-stranded assembly made of stacked guanine quartets. The quartets are stabilized by hydrogen bonds between the bases, and their stacking is further stabilized by monovalent cation coordination in-between the G-quartets. The existence of DNA G-quadruplex structures in cells was revealed thanks to antibodies.^2^ Owing to their prevalence in telomeres and oncogene promoters, G-quadruplexes became attractive targets for anti-cancer therapies.^3, 45–7^ One strategy consists in developing synthetic ligands able to displace the folding equilibria towards the G-quadruplex structure to either hamper telomere maintenance or downregulate oncoproteins. Characterizing the binding affinity, structural selectivity, and G-quadruplex stabilization properties of small molecule ligands is thus of prime interest.

X-ray crystallography^8^ and nuclear magnetic resonance (NMR)^9,10^ are powerful techniques to determine the structure of complexes, but are difficult to apply to polymorphic targets such as the human telomeric sequence (TTAGGG repeats).^11^ To date, six distinct topologies were solved for the target sequences N_i_GGG(TTAGGG)_3_N_j_, depending on the cation and on the choice of overhangs N_i_ and N_j_. The sequence 22AG (see Table 1) forms an antiparallel 3-quartet structure in sodium solution,^12^ a parallel 3-quartet structure in potassium crystals,^13^ and a mixture of at least three different other structures in potassium solutions: the 3-quartet hybrid-1 (solved by NMR for 24TTG^14^ and 23TAG^15^) and hybrid-2 structures^16^ and one antiparallel 2-quartet structure with slipped strands (solved for 22GT^17^). Most structures available for ligand complexes with intramolecular telomeric G-quadruplexes were solved by X-ray crystallography, and the only complexes solved by NMR^16,18,19^ in potassium were cases where the ligand did not significantly affect the topology, i.e. the arrangement of the G-quadruplex stem.

**Table 1.**
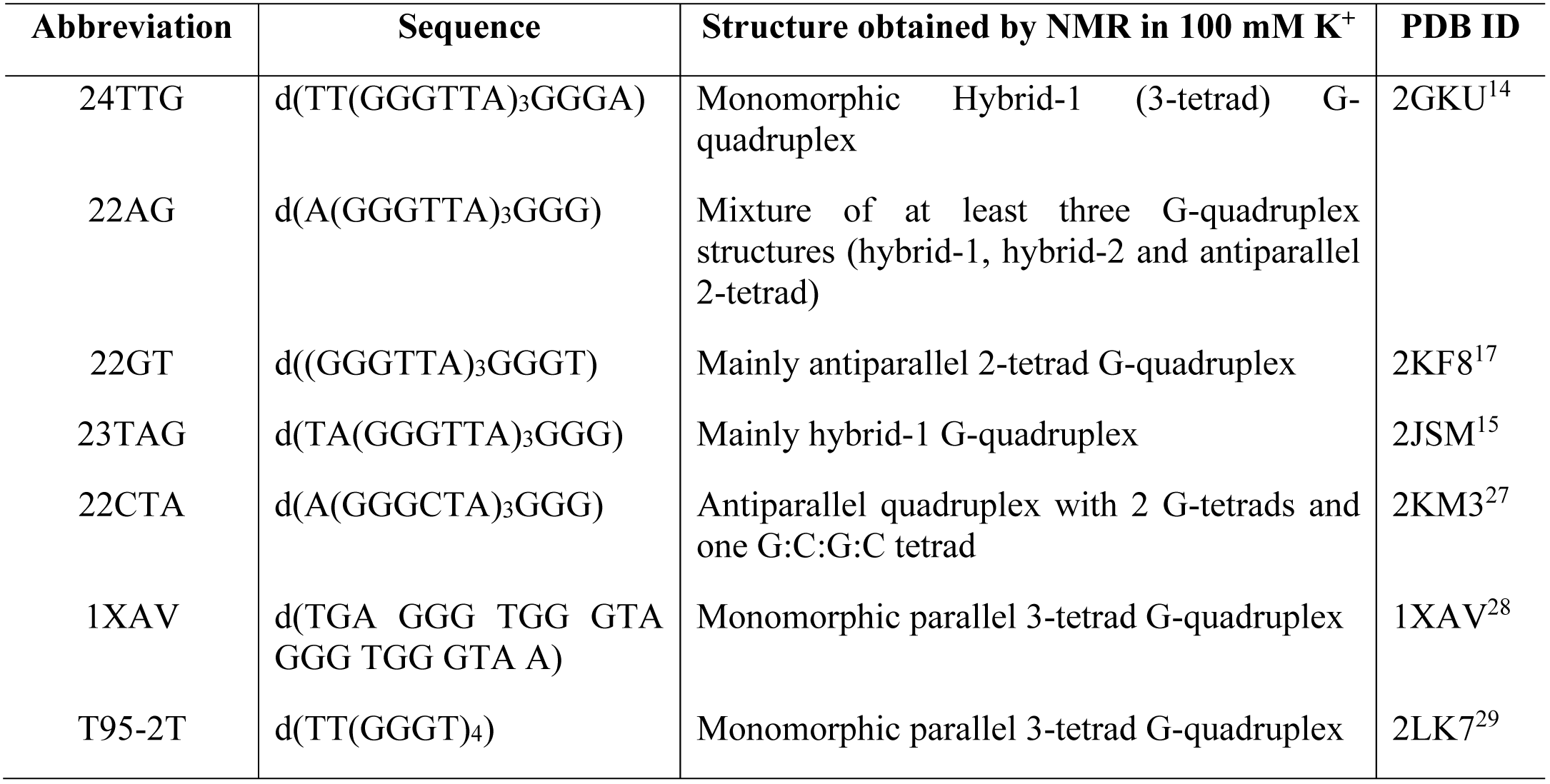
Abbreviations, sequences, structures and PDB codes of the G-quadruplexes studied here.

The scarcity of structural information about ligand binding necessitates to rely heavily on biophysical characterization. Moreover, besides structure, the equilibrium binding constants are the key determinants for the ligand potency, either as a drug or as an artificial probe. Ligand binding constants are determined from isothermal titration experiments, which can be monitored using ligand spectroscopic properties, surface plasmon resonance (SPR),^20^ or calorimetry (ITC).^21^ Another popular method is to monitor a physical property of the folded state as a function of a perturbation such as temperature, and compare the effects in absence and presence of ligand. For G-quadruplexes, thermal denaturation^22^ can be monitored by UV absorption at 295 nm,^23^ by circular dichroism (CD) at wavelengths characteristic of homo-stacking (260 nm) and hetero-stacking (295 nm) of guanines,^24,25^ by Förster resonance energy transfer between chromophores grafted on strand extremities,^22^ by NMR, or by differential scanning calorimetry (DSC).^26^ The temperature at which half of the population is unfolded is called the melting temperature (*T*_m_), and the difference of melting temperature in presence and absence of ligand (Δ*T*_m_) often provides the first ranking when screening ligand libraries for their potential to stabilize G-quadruplexes.

Thermal denaturation data and isothermal titration data do however not always agree, mainly because different states can be populated at different temperatures. In UV-melting or FRET-melting, for example, only one signal is monitored and thus a two-state model often has to be assumed.^30^ Deconvolution of multi-wavelength circular dichroism (CD-melting) signals showed that G-quadruplex melting, without ligand, is not two-state.^31–33^ The same was found with DSC.^34^

With ligands, the situation is expected to become even more complex, because some ligands are able to change the G-quadruplex topology. For example, the very affine and selective telomeric ligand Phen-DC3 conformationally selects a 2-quartet antiparallel structure,^35^ and the Cu(II)-tolylterpyridine selects a 3-quartet antiparallel chair structure upon cooperative binding of two ligands.^36^ These conclusions were based on isothermal titrations monitored by CD and native electrospray mass spectrometry (MS).

The strength of native MS is to unambiguously detect and quantify each of the multiple stoichiometries coexisting in a mixture. In the case of ligand binding to G-quadruplexes, the stoichiometry of each strand:cation:ligand ternary complex (M_n_K_i_L_j_) can be determined unambiguously.^37^ In G-quadruplexes, the number *i* of specifically bound K^+^ ions informs us on the number of stacked G-quartets (*i*+1), and the K^+^ binding is thus a proxy for G-quadruplex folding. Based on K^+^ binding, we recently studied G-quadruplex folding pathways^38^ and demonstrated that 1-K^+^/2-quartet antiparallel G-quadruplexes form at low K^+^ concentration or as short-lived misfolded species. In presence of ligands, the number of specifically bound K^+^ ions indicates whether the complex is unfolded (0 K^+^), has two G-quartets (1 K^+^) or three G-quartets (2 K^+^). Thus, if up to two ligands can bind per G-quadruplex, up to nine states can be readily separated by native MS (Figure 1), and quantified based on the signal intensities^39^ to determine equilibrium binding constants and thus ΔG° of complex formation.

**Figure 1.**
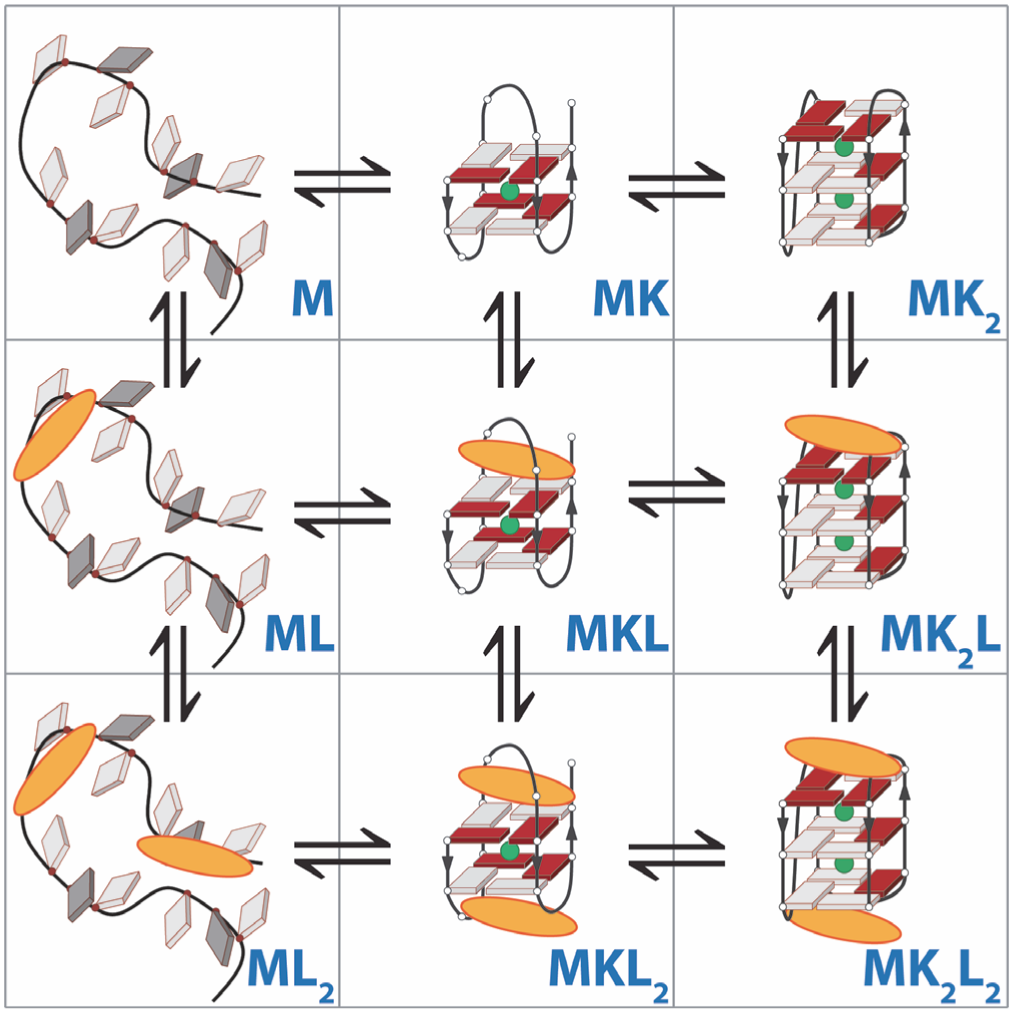
Chemical equilibria for ligand binding to a G-quadruplex forming sequence. Ligand L is in yellow, and potassium ions (K) are in green. G-quadruplexes require at least one potassium ion to fold.

Here, we examined how these binding constants change with the solution temperature. To do so, we built a temperature-variable nanoelectrospray source, inspired by previous designs.^40–43^ The nanospray emitter is embedded inside a temperature-controlled copper block (Figure S 1). MS-melting experiments, i.e. the quantitation of each stoichiometry as a function of the solution temperature, allows a thorough thermodynamic characterization of binding enthalpies (ΔH°) and entropies (ΔS°), and we will show how these thermodynamic parameters can be interpreted in terms of structure.

## RESULTS

### Thermal denaturation of G-quadruplexes

First, we monitored the thermal denaturation of human telomeric G-quadruplexes, without ligands, using CD and native MS. The sequences studied here are listed in Table 1. The CD-melting of 24TTG in 100 mM trimethylammonium acetate (TMAA) and 1 mM KCl is shown in Figure 2. The apparent melting temperature depends on the chosen wavelength. At 290 nm (Figure 2C, black symbols), a wavelength monitoring alternate (*anti*-*syn* and *syn*-*anti*) stacking between adjacent G-quartets, the melting curve is sigmoidal with a *T*_m_ = 40 °C. The inflexion point corresponds to a drop of signal by one half. In contrast, at 260 nm (Figure 2D, black symbols), a wavelength characteristic of homo-stacking (*anti*-*anti*), the CD signal is constant up to ~40 °C, then decreases monotonically, preventing one to accurately determine a melting temperature. The temperature of signal half-drop (*T*_1/2_) is 54 °C. Clearly, more than two states are populated during the thermal unfolding, as also suggested by the absence of isodichroic point in Figure 2A. The presence of intermediates, assigned to parallel structures based on the CD signal at 260 nm, was previously reported by Chaires and coworkers.^32^

**Figure 2.**
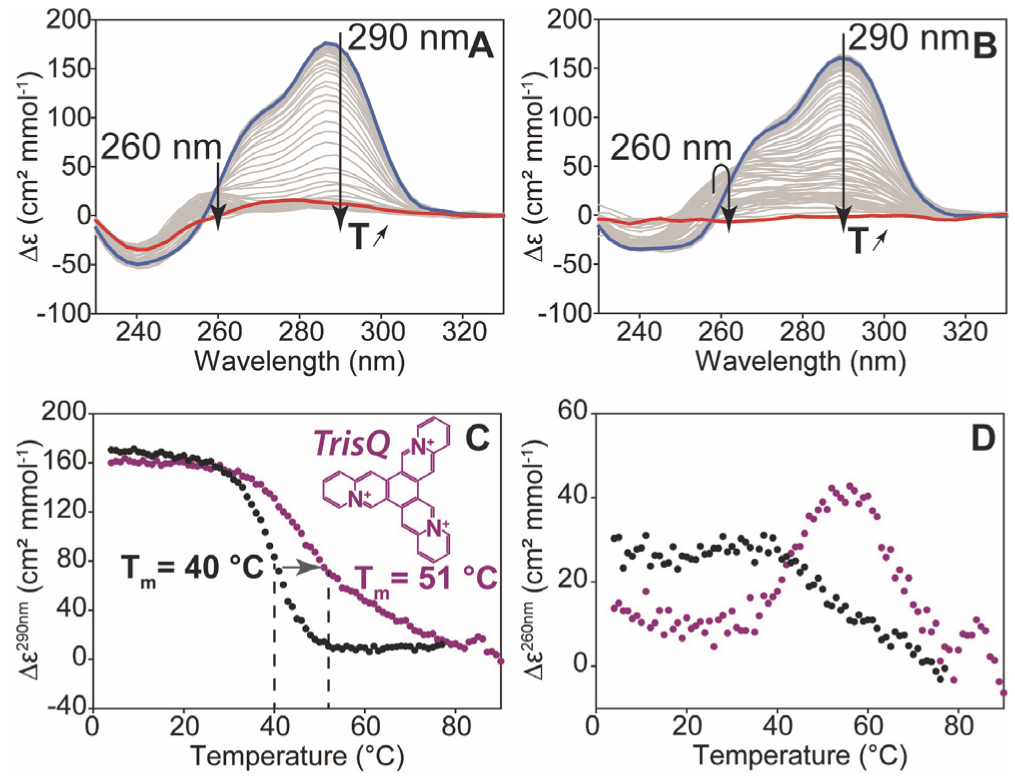
CD-melting of 24TTG (d(TT(GGGTTA)_3_GGGA)) in 100 mM TMAA and 1 mM KCl, without ligand (A) and with one equivalent TrisQ (B). A-B: full CD traces from 4 °C (blue) to 78°C (A) or 90°C (B) (red). Traces at intermediate temperatures are in grey. C-D: Extracted molar circular dichroic signal as a function of temperature for 24TTG (d(TT(GGGTTA)_3_GGGA)) alone (black) or with one equivalent TrisQ (purple) at 290 nm (C) and 260 nm (D).

We then monitored the thermal denaturation by native mass spectrometry (Figure 3). At 26 °C in 1 mM KCl, 24TTG (Figure 3A and Figure S 3) binds exclusively two specific K^+^ ions, indicating that a 3-quartet G-quadruplex is fully formed. When the temperature is increased, the MK_2_ complex disappears and the potassium-free, unfolded state (M) appears. A peak corresponding to the MK stoichiometry is visible, but after correcting for the intensity of nonspecific adducts on M (see experimental details), the abundance of *specific* MK complex is negligible at all temperatures. The abundances of specific M and MK_2_ complexes are then calculated from the intensities, and plotted as a function of temperature (Figure 3B). The melting temperature (50% of unfolded form M) is 46 °C. Only two states can be distinguished based on stoichiometry, suggesting that the additional minor states suggested by CD have a potassium binding stoichiometry of either 0 K^+^ or 2 K^+^. Ion mobility separation did not reveal any significant change of the conformational population under each stoichiometry (Figure S 2).

**Figure 3.**
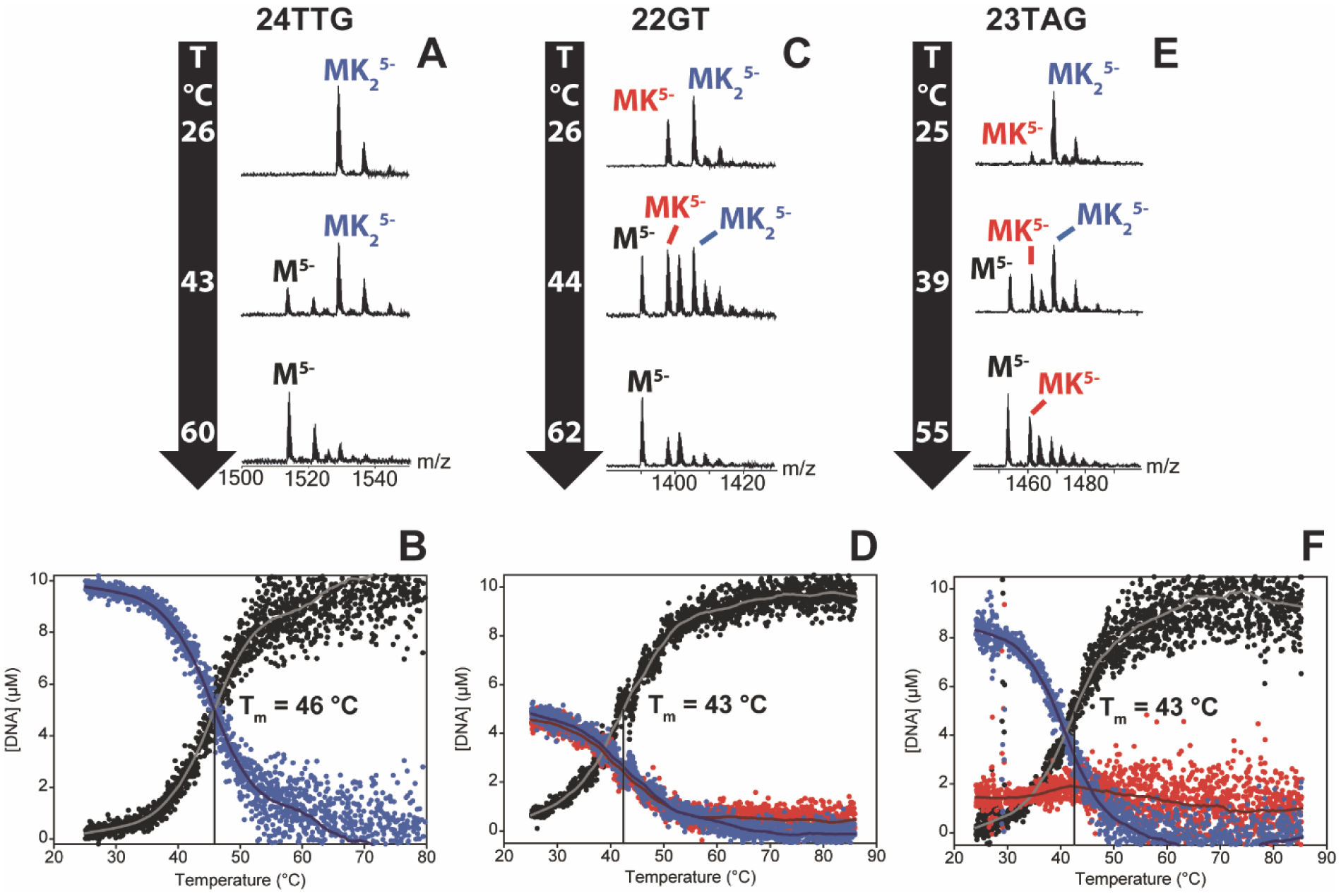
Thermal denaturation experiment monitored using MS of 10 μM 24TTG (A-B), 22GT (C-D) and 23TAG (E-F) in 100 mM TMAA and 1 mM KCl. A, C and E are representative mass spectra of the melting experiment, obtained at the temperature indicated in the black arrows. B, D and F are obtained after quantifying each of the stoichiometry as a function of time and after subtracting the nonspecific adducts contributions (see experimental section).

We also performed MS-melting experiments on telomeric sequences that bind potassium ions less cooperatively than 24TTG, i.e. for which the 1-K^+^ complex is observable. The sequence 22GT forms both MK and MK_2_ complexes at room temperature in 1 mM KCl. MK and MK_2_ disappear concertedly upon increasing temperature (Figure 3B and Figure S 4). The MS-melting temperature is the same for both stoichiometries (43 °C) and is also very close to the one obtained by CD-melting.^35^ The sequence 23TAG folds into a hybrid-1 G-quadruplex very similar to 24TTG in 100 mM K^+^, but can also form a 1-K^+^/2-quartet antiparallel structure at lower KCl concentrations.^38^ At room temperature in 1 mM KCl, the MK_2_ complex predominates (Figure 3C and Figure S 5). When the temperature is increased, the MK_2_ complex disappears, but some MK complex, even after subtraction of the contribution of nonspecific adducts to the peak area, remains present and even slightly increases in abundance above 35 °C. The MK complex thus qualifies as a thermal denaturation intermediate. The melting temperature (50% of M) is 43 °C. The comparison between 23TAG and 22GT illustrate that very different denaturation mechanisms can be hidden behind identical *T*_m_ values.

Results obtained with other human telomeric G-quadruplex forming sequences (22AG and 22CTA) and for two parallel-stranded 3-quartet G-quadruplexes (the c-myc promoter 1XAV and the synthetic sequence T95-2T) are also shown in supporting information (respectively in Figure S 6, Figure S 7, Figure S 8 and Figure S 9). 22AG behaves similarly to 22GT (see more in the discussion section), 22CTA binds only 1 K^+^ ion specifically, and the parallel sequences bind exclusively, and cooperatively, 2 K^+^ ions. No intermediate of peculiar K^+^ binding stoichiometry was observed in those cases.

### Thermodynamics of K^+^ binding to G-quadruplexes

The quantification of each specific complex at each temperature from the peak areas allows one to calculate the equilibrium association constants, and thus the Gibbs free energy at each temperature. Then, using Van’t Hoff plots, the enthalpic and entropic contributions can be determined. The values obtained from the MS-melting experiments are reported in Table 2.

**Table 2.** Thermodynamic parameters of potassium binding to G-quadruplex forming sequences.

| | <sup>MS</sup>$T_m$ (50% M) in 1 mM KCl and 100 mM TMAA (°C) | $\Delta G^\circ$ (298K)<br>(kcal/mol) | | $\Delta H^\circ$<br>(kcal/mol) | | $\Delta S^\circ$<br>(cal/(mol.K)) | | -298 $\Delta S^\circ$<br>(kcal/mol) | |
| --- | --- | --- | --- | --- | --- | --- | --- | --- | --- |
| | | M+K⇌MK | M+2K⇌MK<sub>2</sub> | M+K⇌MK | M+2K⇌MK<sub>2</sub> | M+K⇌MK | M+2K⇌MK<sub>2</sub> | M+K⇌MK | M+2K⇌MK<sub>2</sub> |
| 1XAV | 60 |  | -14.5 ± 1.5<sup>c</sup> |  | -60.1 ± 0.8<sup>a</sup> |  | -153.1 ± 2.3<sup>a</sup> |  | 45.6 ± 0.7<sup>a</sup> |
| T95-2T | 65 |  | -15.3 ± 1.5<sup>c</sup> |  | -55 ± 0.8<sup>a</sup> |  | -135.2 ± 2.3<sup>a</sup> |  | 40.3 ± 0.7<sup>a</sup> |
| 24TTG | 46 |  | -11.48 ± 0.10<sup>b</sup> |  | -53 ± 3<sup>b</sup> |  | -140 ± 10<sup>b</sup> |  | 42 ± 3<sup>b</sup> |
| 23TAG | 43 | -4.9 ± 1.3<sup>c</sup> | -10.3 ± 2.1<sup>c</sup> | -25.3 ± 0.7<sup>a</sup> | -43 ± 1<sup>a</sup> | -68.3 ± 2.1<sup>a</sup> | -109 ± 4<sup>a</sup> | 20.4 ± 0.6<sup>a</sup> | 32.5 ± 1.1<sup>a</sup> |
| 22GT | 43 | -5.32 ± 0.09<sup>b</sup> | -9.4 ± 0.2<sup>b</sup> | -32 ± 3<sup>b</sup> | -30.5 ± 1.8<sup>b</sup> | -89 ± 9<sup>b</sup> | -71 ± 5<sup>b</sup> | 26 ± 3<sup>b</sup> | 21.1 ± 1.6<sup>b</sup> |
| 22AG | 35 | -5.2 ± 0.6<sup>b</sup> | -9.2 ± 0.7<sup>b</sup> | -28 ± 3<sup>b</sup> | -31 ± 3<sup>b</sup> | -75 ± 9<sup>b</sup> | -72 ± 7<sup>b</sup> | 22 ± 3<sup>b</sup> | 21.4 ± 2.2<sup>b</sup> |
| 22CTA | 34 | -4.7 ± 1.0<sup>c</sup> |  | -21.7 ± 0.5<sup>a</sup> |  | -56.9 ± 1.7<sup>a</sup> |  | 17 ± 0.5<sup>a</sup> |  |

<sup>a</sup> Uncertainty = standard deviation from the fit of the Van’t Hoff plot. <sup>b</sup> Uncertainty = standard deviation of multiple experiments. <sup>c</sup> Uncertainty given by the propagation of the standard deviation from the fit of the Van’t Hoff plot.

G-quadruplex formation is enthalpically favorable (Δ*H°*_assn_ < 0) and entropically unfavorable (Δ*S°*_assn_ < 0) in all cases.^44–46^ This is in line with what happens to the DNA sequence: the entropically unfavorable stiffening of the structure is compensated by the enthalpically favorable hydrogen bond formation, stacking interactions, and cation coordination. These effects thus seem to predominate compared to the entropically-favorable solvent release from the solvation shell of potassium and/or from the DNA strand upon G-quadruplex formation (the latter being less hydrated than the former^47,48^).

In Table 2, the sequences are ordered from the most enthalpically favored MK_2_ complex to the least favored (22CTA, in which MK_2_ is not formed). The free energy (Δ*G*°_*assn*_) at 298 K follows the same order. The melting temperatures follow the same trend, except for 1XAV and T92-2T which are reversed. This means that the entropic contributions differ enough so that the stability order is reversed at 298 K compared to around their melting temperature (335-340 K).

The parallel G-quadruplexes 1XAV and T95-2T are the most favored, followed very closely by the hybrid-1 G-quadruplex formed by 24TTG, which is the most monomorphic telomeric G-quadruplex sequence variant. 23TAG, which forms predominantly the same hybrid-1 structure, but also a 2-quartet structure MK, comes next. Note that here, the MK_2_ peak intensity may be the sum of two states: the 3-quartet hybrid-1 structure, and the 2-quartet antiparallel structure with one additional potassium ion specifically bound between a G-triplet and a G-quartet. The latter is the major structure of the MK_2_ complex of 22GT.^17^ The thermodynamic profile of the MK_2_ complex of 22AG is very to that of 22GT. Finally, the thermodynamic parameters for 1 K^+^ binding to 22CTA, forming exclusively a 2-quartet structure with no additional specific K^+^ binding site, closely resemble those of the MK complex of 23TAG. The structural interpretation of these thermodynamic parameters will be further addressed in the discussion section.

### Intermediates in the thermal denaturation of DNA G-quadruplexes with ligands

The CD-melting of 24TTG in the presence of one equivalent of TrisQ, a high affinity G-quadruplex ligand,^49^ is shown in Figure 2B and Figure S 10). At 290 nm (Figure 2C, purple symbols), the apparent melting temperature is 51 °C, so the Δ*T*_m_ induced by the ligand is 11°C. Yet the shape of the melting curve is affected by the presence of the ligand, suggesting the presence of other intermediates and/or a drastic change of the thermodynamic parameters of the unfolding of the complex. Furthermore, other wavelengths give different melting curves (see Figure S 10A) with different melting temperatures. 260 nm is an extreme case: in the presence of TrisQ (Figure 2D, purple symbols), the CD-melting curve is not sigmoidal and no *T_m_* can decently be defined. With the ligand Phen-DC3, the results are highly wavelength-dependent as well (Figure S 10B).

MS-melting helps to disentangle the different states based on their stoichiometry. Figure 4 and Figure S 11 show the MS-melting of 24TTG with one equivalent of TrisQ. At room temperature, the ligand binds by forming mainly a MK_2_L complex. The 2:1 (ligand:quadruplex) complex is absent, although the hybrid-1 structure has two potential end-stacking sites. The low-temperature CD spectrum does not indicate a major rearrangement of 24TTG upon TrisQ binding. This means that the two end-stacking sites are not equivalent.^39^ In addition to the MK_2_L complex, a minor MKL complex is also present. TrisQ is thus also able to bind to a 2-quartet structural variant and partly displace the initial population (major hybrid-1) towards that structure.

**Figure 4.**
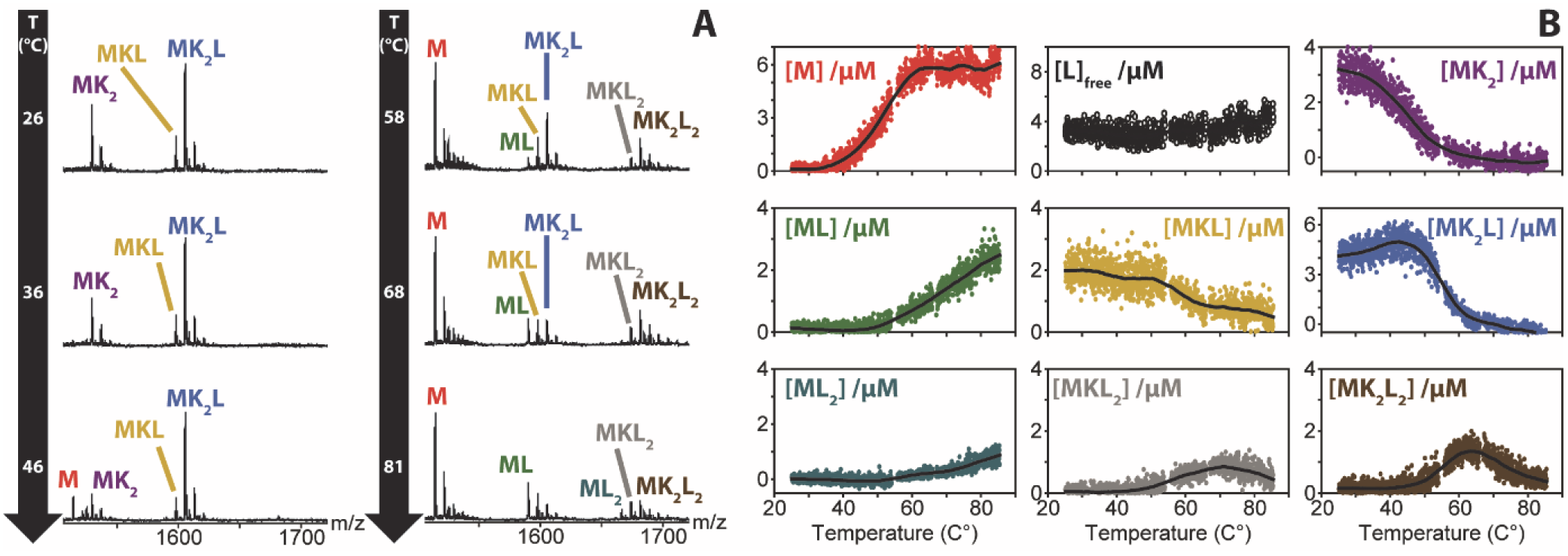
MS-melting of 10 μM 24TTG with 10 μM TrisQ in 100 mM TMAA and 1 mM KCl. A: mass spectra obtained at various temperatures (Zooms on the 5-charge state). B: Quantification of each state distinguished based on its stoichiometry. The concentration of free ligand is obtained by difference.

Upon temperature increase, up to eight states were observed. The sequence of events upon heating is as follows:

1) The MK_2_ state (3-tetrad G-quadruplex alone) melts at a similar temperature as without ligand (~46 °C). This is expected if the experiment reflects equilibria; the only reason for which the structure of the free G-quadruplex would change in presence of ligand would be a slow rearrangement following ligand unbinding. In the range 25-40 °C, the ratio between MK_2_L and MK_2_ increases when the temperature increases, indicating that the 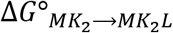 decreases (ligand binding to MK_2_ is increasingly favored).
2) The melting temperature of the main complex (MK_2_L), defined as the temperature at which half MK_2_L disappears, is 55 °C. This temperature is in the same range than—but not identical to—the CD-melting *T*_m_ deduced at 290 nm (51 °C).
3) Surprisingly, new states appear after the denaturation of MK_2_L, notably a 3-quartet complex with two bound ligands: MK_2_L_2_. This counterintuitive result can be explained as follows. At room temperature, the ligand:quadruplex molar ratio is 1:1. However, upon melting of the free G-quadruplex, the ligand:quadruplex molar ratio increases, and TrisQ can thus populate the second binding site of the remaining G-quadruplexes. The formation of new complexes as the temperature increases explains why the spectroscopic melting curves are not sigmoidal. It is also possible that some of the high-temperature complexes have a different structure than the room temperature ones. Intriguingly, the MK_2_L_2_ complex appears at temperatures at which the CD signal at 260 nm increases (Figure 2D), and thus the MK_2_L_2_ complex may contain a higher degree of homo-stacking (typical of parallel-stranded structures). Experiments with T95-2T shows that TrisQ binds to this parallel G-quadruplex with very high affinity and up to 2:1 stoichiometry (Figure S 12).
4) The MK_2_L_2_ complex persists up to much higher temperatures than the MK_2_ and MK_2_L complexes. At the highest temperatures, the MK_2_L_2_ state disappears. Only unfolded species (without K^+^) are formed, and TrisQ ligands are able to bind to these single strands (at least at high temperature).

MS-melting was repeated with three equivalents of TrisQ and gave similar results (Figure S 13), except that up to two TrisQ ligands can bind to the G-quadruplex at room temperature. With 22AG and 1 equivalent TrisQ (Figure S 14), the main difference is the absence of 2:1 ligand:quadruplex stoichiometry at all temperatures. A minor ML_2_ complex was observed only with the unfolded strand.

Other ligands displayed different states upon thermal denaturation, as illustrated in Figure 5 At room temperature, Phen-DC3 binds to the 2-quartet antiparallel form of telomeric G-quadruplexes, predominantly with the MKL stoichiometry when there is one equivalent ligand. Similarly to TrisQ, a 2:1 (ligand:quadruplex) complex MKL_2_ forms above the melting temperature of the free G-quadruplex, for both 24TTG (Figure 5A and Figure S 15) and 22AG (Figure S 16).

**Figure 5.**
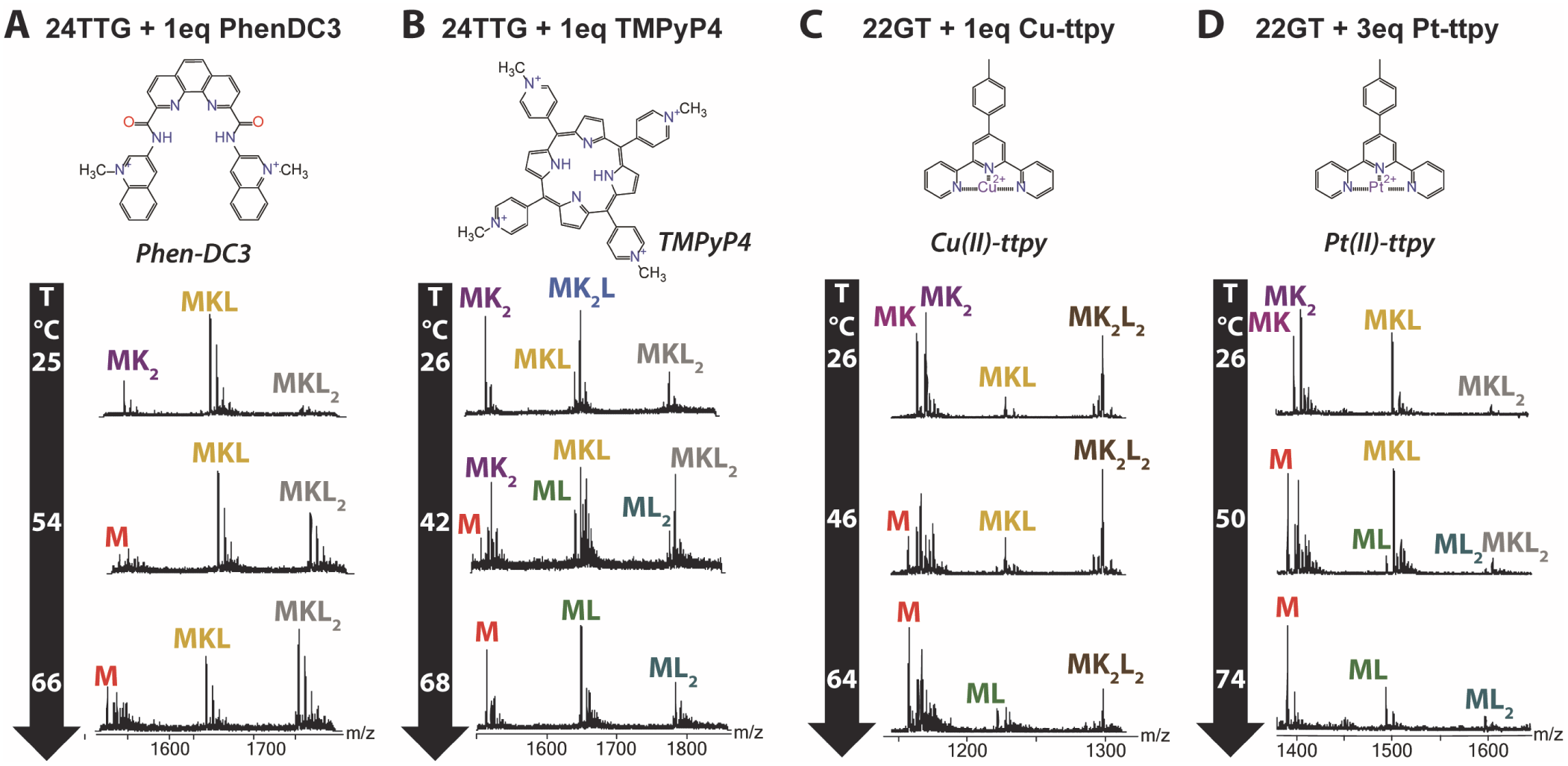
Mass spectra at different temperatures of 10 μM of human telomeric DNA sequence (24TTG, A and B; 22GT, C and D) in the presence of one or three equivalent(s) A, Phen-DC3, B, TMPyP4, C, Cu-ttpy and D, Pt-ttpy.

At room temperature, TMPyP4 binds to 24TTG with a similar profile as TrisQ does (Figure 5B and Figure S 17). However, the states appearing upon melting differ. Upon G-quadruplex melting (indicated by potassium release), the ligand binds to the unfolded form, so the apparent ligand:strand stoichiometries and abundances remain relatively constant. This highlights the poor selectivity of TMPyP4 for the G-quadruplex structures.

We also studied the thermal denaturation of 22GT in presence of two different metal complexes, Cu-tolyltertpyridine (Cu-ttpy) and Pt-tolyltertpyridine (Pt-ttpy) (Figure 5C-D and Figure S 18, Figure S 19 and Figure S 20). Cu-ttpy binds non-covalently and cooperatively to human telomeric G-quadruplexes, and the ligand-bound form is antiparallel with three quartets.^36^ At room temperature, the main complex is MK2L2. MKL is also present, but minor. In contrast, Pt-ttpy binds covalently to adenines in the loops of telomeric sequences.^50,51^ With 22GT, the main complex is MKL. Based on FRET-melting studies in potassium,^50^ Cu-ttpy had been discarded as a poor ligand, whereas Pt-ttpy retained attention owing to its higher Δ*T*_m_. Yet, at 26 °C, the fraction of free G-quadruplex is identical with either 1 equivalent Cu-ttpy (Figure 5C and Figure S 18 and Figure S 19) or 3 equivalents Pt-ttpy (Figure 5D, Figure S 20). This illustrates a case where the ^FRET^Δ*T*_m_ values are a poor indicator of ligand affinity, and led to an erroneous ranking. The reason lies in the very different states involved. For Cu-ttpy, the MK_2_L_2_ species melts at a slightly higher temperature than the ligand-free G-quadruplex. Cu-ttpy also binds noncovalently to the unfolded form at high temperature. In contrast, because Pt-ttpy binds covalently,^50,51^ the complex can be considered a modified oligonucleotide, which unfolds (loses K^+^) at a higher temperature (56 °C) than the unmodified 22GT (43 °C).

### Thermodynamics of ligand binding to G-quadruplexes

In the same way as for the G-quadruplexes alone, we quantified each stoichiometric state along the MS-melting experiments. The free ligand and free potassium ion concentrations were calculated by difference, from the mass balance equation. Knowing all concentrations gives access to the equilibrium binding constants and Δ*G*° of each individual reaction *M + i K^+^ + j L* ⇌ *MK_i_L_j_* at each temperature. When the reactant M and product MK_i_L_j_ were not observed simultaneously at a given temperature, the values were obtained from coupled equilibria. A Van’t Hoff analysis then gives access to Δ*H*° and Δ*S*° of these reactions. The concentration plots and Van’t Hoff plots are all shown in supporting information Figure S 11-Figure S 20. The values of the thermodynamic parameters are gathered in Table S 1. Figure 6 summarizes the key results for discussion, in the form of checkerboards of ΔΔ*G* °, ΔΔ*H*° and ΔΔ*S*° values compared to the predominant G-quadruplex state without ligand (MK_2_ for 24TTG, MK for 22AG). This allows one to visualize the cost or gain upon adding/removing each ligand (moving down/up on the checkerboard) or adding/removing each K^+^ ion (moving right/left on the checkerboard) on the strand M.

**Figure 6.**
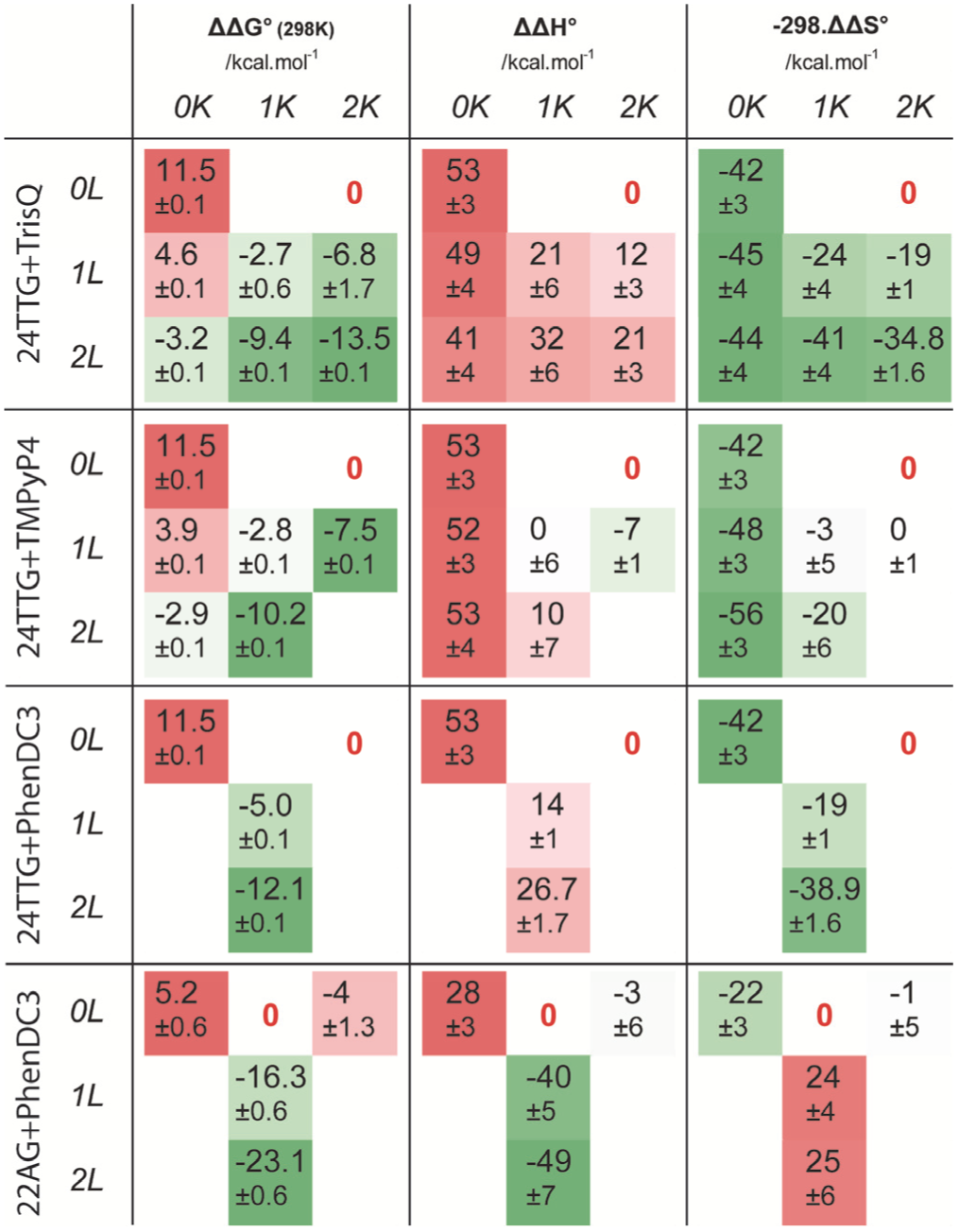
Thermodynamic parameters for potassium cation and ligand binding to G-quadruplexes. ΔΔ*G°*, ΔΔ*H°* and −298.ΔΔ*S°* are relative to the folded G-quadruplex (MK_2_ for 24TTG, MK for 22AG, bold red zero). The values color-coded, indicating how favorable (green) or unfavorable (red) is a given state compared to the free G-quadruplex, should the reaction be carried out at 1 M concentration of each partner. The reported uncertainties are the standard deviations from repeated experiments, considering uncertainty propagation when sequential reactions are used for the Van’t Hoff regressions.

TrisQ binding to 24TTG is entropically favorable. With −298. ΔΔ*S*° around −20 kcal/mol per ligand bound, the system typically regains the entropy it had lost per potassium ion bound. In contrast, TMPyP4 binding to 24TTG is entropically indifferent, and favored only by enthalpy. This indicates a different binding mechanism.

The thermodynamic parsing according to stoichiometry harbors further information related to the conformational changes associated with ligand binding. In particular, MS allows one to isolate the states with only 1 K^+^ ion bound, corresponding to 2-quartet structures, and get individual thermodynamic parameters for that structure. For example, when one Phen-DC3 binds to 24TTG, it ejects one K^+^. This binding, which is accompanied by a major topology rearrangement, is characterized by a favorable entropic contribution (−298. Δ*S*°_MK →MKL_ = −19 *kcal*/*mol*), in line with the liberation of one K^+^ and one G-quartet. In contrast, the binding of one Phen-DC3 to 22AG has a totally different thermodynamic profile: it is enthalpically favored (Δ*H* °_MK→MKL_ = −40 *kcal/mol*) and entropically unfavored (-TΔ*S*°_MK→MKL_ = 24 *kcal/mol*), suggesting either another binding mode to this G-quadruplex, or that the starting G-quadruplex has a very different conformation. This will be further discussed below.

## DISCUSSION

### Uncertainties in the MS-melting method, and comparison with traditional UV-melting, CD-melting or FRET-melting

The contribution of random fluctuations to uncertainty was estimated from replicate experiments. For example, for 24TTG in 1 mM KCl (Table S 2), the standard deviation on the melting temperature is 1 °C, which is similar to UV-melting or CD-melting. One significant source of uncertainty comes from a possible difference between the temperature of the emitter tip, protruding 0.5-1 mm from the copper bloc, and that of the copper block on which the temperature is monitored. We performed Comsol Multiphysics simulations with the block heated from 25 to 70 °C in room temperature (25°C) air (Figure S 21), and found that the temperature of the tip is lower than the temperature of the copper bloc by 0 to −3 °C.

The standard deviation on thermodynamic parameters from replicate experiments was ~5-12 % for Δ*H*° and Δ*S*°, from the Van’t Hoff plots. The uncertainty on Δ*G*298° depends on how far the melting temperature is from 298 K. To verify whether the Δ*H*° and Δ*S*° values obtained by MS-melting are reasonable, we also studied two well-characterized complexes: Hoechst 33258^52^ and Amsacrine^53^ (Scheme S 1) bound to the Dickerson-Drew dodecamer (dCGCGAATTCGCG)_2_. The difference in binding modes (grove binding versus intercalation, respectively) strongly affects the thermodynamic parameters. Chaires showed that the binding of Hoechst 33258 is entropically favored,^54^ and Graves reported that the binding of Amsacrine is enthalpically driven.^53^ Our results (supporting information Figure S 22 and Figure S 23) show that the values of Δ*G*° agree, despite the slight differences in buffer. The values of Δ*H*° and Δ*S*° show more discrepancies, but are always of the same sign. We will thus base our discussion of MS-melting results on the sign and magnitude of the thermodynamic parameters rather than on the absolute values.

A significant source of uncertainty, compared to calorimetric methods, is the use of Van’t Hoff plot, with the underlying assumption that the enthalpy is independent on the temperature. Calorimetry showed that this is usually not the case for G-quadruplexes.^34,55^ Not taking into account the heat capacity could give errors up to 20%.^56^ One possible solution is to perform several MS-melting experiments at different K^+^ and/or ligand concentrations, so as to shift the melting temperature significantly, and thus obtain thermodynamic parameters at different temperatures. We explored this approach with T95-2T at 100 μM and 1 mM KCl (Figure S 24). The difference in Δ*H*_assn_° obtained around 45°C and 65°C is 13%. The Δ*C_P_*_,*assn*_° = −358 *cal K*^−1^ *mol*^−1^ is in very good agreement with values obtained for other G-quadruplexes,^55^ so this approach will be worth exploiting in future experimental designs.

However, compared to all spectroscopic and calorimetric techniques, the significant advantage of MS-melting is the direct readout of the stoichiometry of each state. Consequently, the chemical equilibria for which one extracts the thermodynamic parameters are unambiguously defined. Importantly, multiple equilibria can be tracked simultaneously, whereas other techniques, including calorimetry, would struggle to disentangle the states or to interpret the chemical nature of the states extracted by singular value decomposition.^32^ Finally, defining baselines for each state is not difficult when treating MS-melting data, which is also a significant advantage compared spectroscopy-melting data.^22,30^

### The driving forces for potassium-induced G-quadruplex folding

Using mass spectrometry, we distinguish G-quadruplex folded states via potassium binding, i.e. a multimolecular reaction:

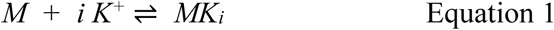

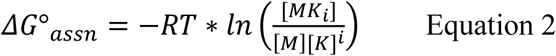

One should be cautious if the MS-melting results are compared with literature in which the folding is defined by a pseudo-unimolecular folding reaction:

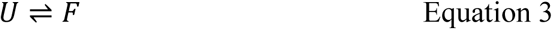

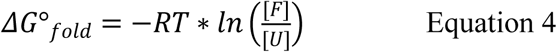
is a *conditional* equilibrium binding constant (i.e., it depends on the experimental conditions, here most notably on the KCl concentration), and Δ*G*°_*fold*_ is related to Δ*G*°_*assn*_ by:

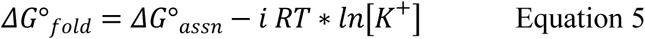

When discussing literature values one must be critical about how the stoichiometries were assigned. In MS, the stoichiometries are unambiguous and each stoichiometric state is quantified. With traditional methods, the determination of *i* is more indirect (one may find non-integer numbers) and this is a possible source of discrepancy. Also, differences can come from other components of the solutions, most notably the ionic strength, which may influence the activity coefficients of DNA forms and of cations. Both in our work and in the G-quadruplex literature, the analytical concentrations are used as if they were equal to activities. Finally, the state partitioning in MS is defined by the stoichiometry, whereas spectroscopic techniques (NMR) or data treatment techniques (SVD) may partition the ensemble into very different groups. For example, while the specific MK stoichiometry is a unique signature for a 2-quartet state (only antiparallel structures are known to date), the MK2 complex can be either parallel, antiparallel or hybrid 3-quartet, or the two-quartet structure with one extra cation coordinated between a quartet and a triplet.

However, the thermodynamics of MK_2_ formation will differ depending on its structure. Figure 7 shows the thermodynamic parameters per specific cation bound for four telomeric sequence variants. For 24TTG, no specific MK complex is detected, and the energy per cation is calculated as half the energy of specific binding of the two cations. We verified by ^1^H NMR that, in our 1-mM KCl conditions, 24TTG forms a hybrid-1 structure and 22GT forms an antiparallel 2-quartet structure, like in 100 mM KCl (supporting information Figure S 25). 23TAG and 22AG show both the MK and the MK_2_ stoichiometry. However, the thermodynamics of cation binding are very different.

**Figure 7.**
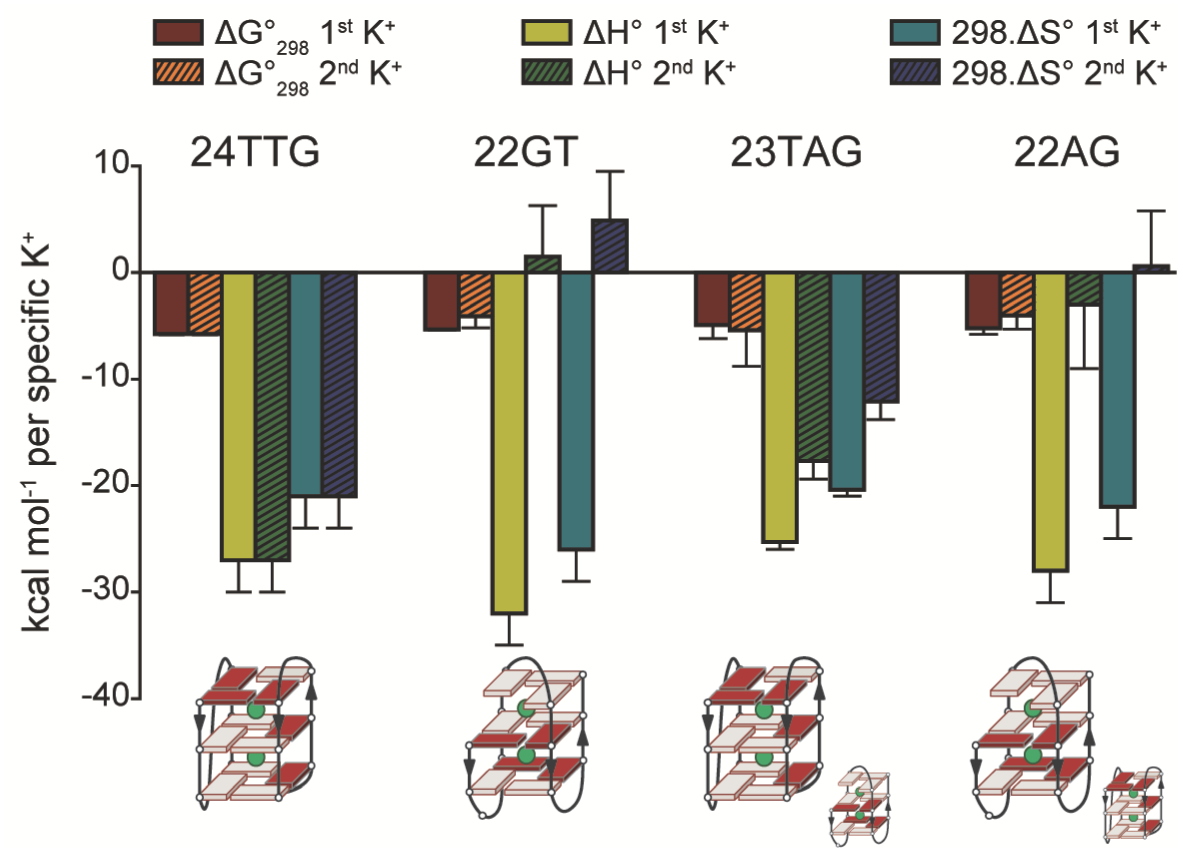
Δ*G*°, Δ*H*° and *T*Δ*S*° per cation for four different telomeric sequence variants. Bottom: structure of the MK_2_ complex, based on NMR for 24TTG and 22GT, and deduced predominant, based on thermodynamic signature, for 23TAG and 22AG.

The similar thermodynamic profiles of potassium binding to 22AG and 22GT suggest that the G-quadruplexes formed by 22AG and 22GT are very similar. For 22GT, which folds into an antiparallel 2-quartet with a triplet of guanines at up to 100 mM K^+^,^17^ the first K^+^ is expected to bind between the two quartets, and the second K^+^ could bind in-between one quartet and the triplet of guanines, without altering the MK structure. Our results suggest that 22AG also forms a similarly complex ensemble of structures in the mM K^+^ concentration range; the ^1^H NMR spectrum shows multiple coexisting species (Figure S 25). When compared to the fractions predicted by Boncina *et al*. for states U, I and F, the 2-quartet structure would correspond to the intermediate I. It is likely that, at higher K^+^ concentration, the fraction of truly 3-quartet structures increases. The comparison between 23TAG and 22AG shows that a single thymine on the 5’-end suffices to drive the equilibrium significantly towards the hybrid-1 structure. The potassium binding thermodynamics for the MK_2_ complex of 23TAG is much more similar to that of 24TTG, and the major peaks in the ^1^H NMR spectrum of 23TAG at 1 mM KCl are identical to those found in 100 mM KCl, where a hybrid-1 structure was solved.

The main difference between the different potassium binding modes is that the formation of each quartet-K^+^-quartet units is strongly enthalpically favored and entropically unfavored, whereas the formation of a extra quartet-K^+^-triplet unit is entropically favored. This means that the driving force for proper G-quadruplex folding (G-quartets only) comes from bond formation, and that this requires significantly stiffening the conformation. However, the cation-mediated stacking of a triplet is entropically favorable despite the additional conformational restrictions, and the main contribution may thus be water release.

Also, a positive *T*Δ*S*° contribution for triplet formation means that triplet formation is favored when the temperature is increased. This is why the population of the “misfolded” (two quartets instead of three) structures can increase with temperature, as observed for the MK state in 23TAG (Figure 3F) or for the intermediate state (I) defined by *Boncina et al*. Finally, the positive *T*Δ*S*° for potassium-stabilized triplet formation gives some credit to the hypothesis of “G-triplex” formation at intermediate temperatures upon thermal denaturation.^57,58^ Although we do not observe here pure G-triplexes, our results suggest that potassium-stabilized guanine triplets may indeed be favored at higher temperatures than pure quartets, for entropic reasons.

### Definition of the melting temperature (T_m_), and pitfalls in using spectroscopy-based ΔT_m_ values to rank ligand binding affinities

The *T*_m_ is the temperature at which half of the population is unfolded. In MS-melting of G-quadruplexes, the *T*_m_ can be inferred from the concentration of all potassium-free forms (with and without ligand). In MS-melting of intermolecular complexes (e.g., duplexes), the unfolded state corresponds to the single strands. In spectroscopic techniques, the *T*_m_ is often estimated from the median between the high-temperature and low-temperature baselines.^30^ However, when many ensembles are populated as a function of temperature, which often happens in presence of ligands, this simplistic approach fails because the transitions are no longer sigmoidal (the temperature of half-drop of signal must be used instead, and the annotation *T*_1/2_ is then preferred). For example, for the binding of TrisQ to 24TTG in Figure 4, up to eight ensembles are populated. In addition, some ensembles increase, then decrease over the melting experiment. MS-melting offers an unbiased way to determine melting temperatures, because the signal of each folded and unfolded state is quantified simultaneously.

In the G-quadruplex research community, ligand screening often starts with a ranking according to a melting assay. A common way to rank ligand binding affinities is to calculate the Δ*T*_m_ or Δ*T*_1/2_ from experiments with and without ligand. Disentangling the states by native MS reveals the potential pitfalls when screening ligands using Δ*T*_m_ values. For identical Δ*G*°_298_ values, entropically favored ligand binding modes give higher Δ*T*_m_ values than enthalpically favored ligand binding modes. Thus, when ligands having different binding modes are compared within a series, the Δ*T*_m_ ranking is not a good predictor of affinity ranking at other temperatures. MS-melting experiments allow to estimate the equilibrium binding constants (affinities) at each temperature, and are thus key to lift discrepancies between melting experiments and isothermal *K*_D_ determinations. Furthermore, Δ*T*_m_ ranking may have led to favoring ligands having an entropically-favored binding mode compared to ligands with enthalpically-driven binding. The prevalence of melting assays as first screen may have contributed to lower the diversity of ligand binding modes discovered until now.

### Thermodynamic signatures of ligand binding, including ligand-induced conformational changes

Planar aromatic ligands such as TrisQ are expected to stack on external G-quartets.^59,60^ We found that the binding of TrisQ to 24TTG is enthalpically unfavorable. This is compatible with stacking because in the ligand-free form, external G-quartets are often capped by additional base pairs.^19^ Ligand end-stacking disrupts these interactions, resulting in unfavorable enthalpic and favorable entropic contributions. The release of solvation ions/molecules from the ligand and the G-quadruplex can also contribute to the entropy gain.

In line with previous reports based on calorimetry,^61,62^ we found that the binding of Phen-DC3 to 22AG is enthalpically driven. The previous studies suggested that the displacement of water molecules from the binding interface is responsible for the high affinity. However, we found that the thermodynamic signature for the binding of Phen-DC3 to 24TTG is very different compared to 22AG, so the hypothesis of water release as a driving force does not hold. Our interpretation is that the thermodynamic signatures reflect that the starting structures are different. From potassium binding thermodynamics, we concluded above that 22AG folds mainly into 22GT-like structures (2-quartets 1-triplet of guanines antiparallel G-quadruplex) in our experimental conditions, so there is no major topology rearrangement upon binding of Phen-DC3 to 22AG. We propose that the enthalpically favorable and entropically unfavorable character of Phen-DC3 binding to the 2-quartet structure is due to the formation of new hydrogen bonds between the ligand and the guanines which are left free on both ends of the 2-quartet core. In contrast, upon binding of Phen-DC3, 24TTG therefore has to completely rearrange its topology, and this contribution translates into major differences in the ligand binding thermodynamics.

For TMPyP4, *Lah et al*. described that the binding of TMPyP4 to 22AG is mainly driven by entropy.^62^ Despite the fact the CD signals in presence of the ligand decreased significantly, the authors assumed that no particular conformational changes were occurring upon binding of TMPyP4, and interpreted the entropically-driven binding as due to solvent release from the buried hydrophobic surfaces. Our MS-melting results provide a much more complete picture. With the sequence 24TTG, we could separately characterize TMPyP4 binding to the 3-quartet hybrid structure, to the 2-quartet structure, and to the unfolded form. The binding to the hybrid-1 structure is enthalpically favored, the binding of to the 2-quartet structure is slightly entropically favored, and the binding to the unfolded structure is very entropically favored. The binding mode of TMPyP4 to G-quadruplexes has remained controversial: some reports show that it binds and stabilizes G-quadruplexes,^63^ others show that TMPyP4 changes the G-quadruplex structure,^64^ and others show that it unfolds G-quadruplexes.^65–67^ Here we show that all types of binding coexist, and that the distribution of final states depends on the ligand concentration ratio and on the temperature.

## CONCLUSIONS

By analyzing G-quadruplexes formed in the presence of potassium and their complexes with ligands using a temperature-controlled electrospray source, we could, in a single melting experiment, determine the enthalpy, entropy and free energy of formation of each complex based on its stoichiometry. This allowed us to study the thermodynamics of G-quadruplex folding, one potassium ion at a time. Potassium binding between a G-quartet and a triplet has a very different thermodynamic profile than the formation of a quartet‒K^+^‒quartet unit, because it is entropically favored. The cause may be that the entropy of water release upon K^+^ binding is larger than the conformational entropy loss due to the triplet formation. An important consequence is that slipped structures may become favorable as the temperature is increased, below the melting temperature at which the entire G-quadruplex unravels.

In several cases, the mass spectra revealed states that were not suspected based on traditional melting assays, and MS-melting experiments are particularly useful to explain apparent inconsistencies between isothermal and melting assays. The ability to distinguish between unfolded structures (no K^+^ bound), 2-quartet structures (1 K^+^ bound) and 3-quartet structures (2 K^+^ bound) is important, because several popular ligands (360A, Phen-DC3) are capable of inducing major changes of topology towards 2-quartet folds in telomeric sequences. We also observed such complexes with the ligands TrisQ, TMPyP4 and Cu-ttpy.

In the melting of ligand complexes, we also found that, upon unfolding of the main 1:1 (ligand:quadruplex) complex, 2:1 complexes are formed alongside unfolded forms. This comes from a change of stoichiometric ratio between ligand and (remaining) quadruplex as the ligand-free fraction unfolds. Thus, even when a melting experiment is carried out with one equivalent of ligand, the melting of the higher stoichiometries may ultimately determine the magnitude of the Δ*T*_m_.

We also found that when the ligand binding stoichiometries differ from ligand to ligand, or when ligand binding is associated with topology changes, the Δ*T*_m_ values do not reflect the ligand binding constants. The full thermodynamic profiles for ligand binding thus bring important insight into the driving forces for ligand binding. The present study highlighted how the thermodynamics contributions depended on the binding scenarios. For example, ligand end-stacking (TrisQ on 24TTG), releasing both water bound to the ligand and conformational strain in end-stacked bases, is favored entropically, whereas ligand hydrogen bonding (hypothesized for Phen-DC3 on 22AG) translates into more favorable enthalpic contributions. Major topology changes upon ligand binding also translates into specific thermodynamic signatures, which are particularly useful given that no high-resolution structure of ligands inducing changes of topology is currently available.

In the future, the most powerful thermodynamic characterization of nucleic acid/drug melting will come from a combination of techniques. The quantitative parsing among different states based on their stoichiometry is the key advantage of mass spectrometry. The results illustrate the richness of information obtained from MS-melting experiments alone. Combining MS-melting with CD-melting can further parse the states (especially the MK_2_ complexes) according to stacking topology, and differential scanning calorimetry experiments in the same experimental conditions would provide key information on the contribution of solvation effects thanks to Δ*C*°_p_ values. Such full thermodynamic characterization should provide key insight into the driving forces for G-quadruplex cation-induced folding and ligand binding.

## MATERIALS AND METHODS

### Oligonucleotides and ligands

Oligonucleotides were purchased lyophilized and RP-cartridge purified from Eurogentec (Seraing, Belgium). Stock solutions were prepared at 0.80 or 1.00 mM concentration in nuclease-free water (Ambion, Fisher Scientific, Illkirch, France). The concentrations were determined using the Beer-Lambert law by measuring the absorbance at 260 nm. The extinction coefficients were obtained from the IDT website using the Cavaluzzi-Borer correction.^68^ The solutions were then diluted to 200 μM before being prepared into the folding conditions. Trimethylammonium acetate (TMAA, Ultra for UPLC) ammonium acetate (NH_4_OAc, BioUltra for molecular biology, ~5 M in H_2_O) and potassium chloride (>99.999%) were purchased from Sigma-Aldrich (Saint-Quentin Fallavier, France).

The ligands TrisQ and Phen-DC3 were synthesized as described elsewhere^49,69^ and provided by Prof. Teulade-Fichou from Institut Curie, Paris. TrisQ and Phen-DC3 were solubilized at 2 mM in DMSO before being diluted to 500 μM using water. TMPyP4 was purchased from Sigma-Aldrich. The concentration of these three ligands was determined using UV absorbance. The following extinction coefficients and wavelengths were used: for Phen-DC3: ε_320_ = 62400 cm^−1^ M^−1^, for TrisQ ε257 = 70900 cm^−1^ M^−1^ and for TMPyP4 ε_424_ = 226000 cm^−1^ M^−1^. Cu-ttpy and Pt-ttpy were synthesized in house using previously established protocols.^36,50^ Their concentrations were determined from careful weighing. Amsacrine hydrochloride was obtained from Sigma and the bisbenzimide Hoechst 33258 from Sigma-Aldrich (Saint-Quentin Fallavier, France). Table 1 lists the G-quadruplex forming sequences used in this article. Moreover, we used the control sequences 21nonG4 (d(GGGATGCGACGAGAGGACGGG)), 22nonG4 (d(GGGATGCGACAGAGAGGACGGG)) and the self-complementary duplex-forming sequences DK33 (d(CGTAAATTTACG),) DK66 (d(CGCGAATTCGCG)) and DK100 (d(CGCGGGCCCGCG)) for control experiments including the tests with minor groove binders and intercalators.

### Native mass spectrometry

Melting experiments were performed on an Agilent 6560 IMS-Q-TOF (Agilent Technologies, Santa Clara, CA, USA), using a home built source described in the next section. The experiments were performed in negative mode. The instrument allows recording the drift time distribution of each *m*/*z* species, which helps to discern eventual conformational ensembles. One MS spectrum is summed every ~1 s. The borosilicate nanospray emitters were purchased from ThermoFisher (ref. ES388) and opened manually. The capillary voltage was set between 1.0 and 1.5 kV. The spray capillary is grounded and the source entrance voltage is set to high voltage. A backpressure of 0.1 to 1 bar was applied to ensure a continuous production of ions during the entire melting experiment. The gas in the ion mobility cell of the instrument was helium (Alphagaz 2 grade from Air Liquide, H_2_O < 3 ppm, C_n_H_m_ < 0.5 ppm, O_2_ < 2 ppm). The pumping system in the source region was modified using a second Triscroll TS800 pump (Agilent Technologies).^70^ Some experiments (sequence T95-2T) were reproduced on a Synapt G2S (Waters, Manchester, UK), using similar conditions. In this instrument, the cone is grounded and the source is connected to the high voltage.

### Home-built temperature-controlled nanoelectrospray source, TCnESI

The TCnESI source is a classical nESI source in which the entire emitter is embedded in a copper block (Typically, 0.5-1 mm of the emitter protrudes outside of the block) (Figure S 1). The copper piece was made by the ETH workshop. This block is heated or cooled by a Peltier element purchased from RS-online (Adaptive ET-127-10-13-RS, 15.7 V, 37,9 W). Its power is supplied by a Kikusui PMX18-5A power supply (Kikusui Electonics Corp., Japan) which is remote-controlled by a LabView software (National Instruments). The temperature of the copper block was measured using a bolt mount thermistor sensor (5000 Ohms, ON-950-44005, Omega, standard accuracy of 0.2 °C from 0 to 70 °C) plugged into a Keithley 2701E multimeter (Equipements Scientifiques, Garchies, France), itself connected to the LabView software. The LabView code is home-made and consists in a loop adjusting the temperature.

For the melting experiments, a ramp was programmed to scan from 25 to 85 or 95 °C at 2 °C/min. This rather fast ramp maximized the chances of recording the entire melting in a single experiment without losing the spray signal. Typically, a single nanospray emitter, in our conditions, lasts for 30 to 60 minutes. Note that similar ramp speeds are customary in FRET-melting experiments, wherein the small sample wells respond quickly to temperature changes.^32,71^ To check whether this scan speed could affect the results or not, we performed the melting of the same complex (the DK66 duplex in 100 mM NH_4_OAc) using 2 and 1 C°/min and found very similar results (T_m_ = 53.6 and 53.9 °C, supplementary Figure S 26). As an additional control, in supplementary Figure S 27, we show the thermal denaturation of three auto-complementary DNA duplexes containing increasing number of G≡C base pairs (DK33, 33% G≡C, DK66, 66% G≡C and DK100, 100% G≡C) monitored using mass spectrometry. At low temperature, the duplexes (M_2_) are formed in 100 mM NH_4_OAc. When the temperature is increased, the duplexes dissociate into monomers. At high temperature, only the single strand can be observed. The quantification reveals the evolution of the two species as a function of temperature. The curves were fitted using sigmoid functions and the melting temperature (as defined by the inflexion point) determined. As expected due to the increasing number of hydrogen bounds between the two strands, the melting temperature is shifted towards higher temperatures with increasing number of G≡C base pairs (T_m_(DK33) = 43.5 °C < T_m_(DK66) = 53.6 °C < T_m_(DK100) = 64.6 °C).

### Circular dichroism

A Jasco J-815 spectrophotometer was used to perform the CD experiments. A quartz cell of 2 mm path length and 10 mm width was used. The temperature ramp was set to 0.4 °C/min from 4 to 90 °C. The temperature ramp is halted during the acquisition of each spectrum, which consists of two scans acquired at 100 nm/min. A linear baseline was subtracted from each spectrum using the mean value between 320 and 350 nm. CD data were converted to molar circular-dichroic absorption (Δ*ε*) based on DNA total concentration (*C*) using the following equation,

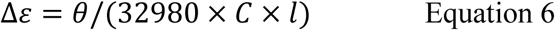
where *θ* is the ellipticity in mdeg and *l* the length of the cell in cm (0.2 cm).

### Mass spectrometry data processing

For the duplexes, the peaks corresponding to the monomer and to the dimer were integrated as a function of temperature. The charge states were summed and normalized to the total concentration. If the peaks overlap in mass (for example a monomer M^3-^ and a dimer 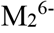), ion mobility is used to separate the peaks (supplementary Figure S 28). We considered all response factors equal. The difference between the duplex melting temperatures obtained by MS and by other methods could come in part from this assumption.

For the G-quadruplexes, the unfolding was monitored by a change in the number of bound K^+^. We found that, upon temperature increase, the shift in charge state is very small and not due to the thermal denaturation of the complexes because similar shifts are observed for non-G-quadruplex forming sequences (Figure S 29 and Figure S 30).

The peaks corresponding to the different K^+^ and L stoichiometries were integrated as a function of temperature. From the relative abundances, the DNA concentrations are obtained and the free K^+^ or ligand concentrations can be obtained by difference. For each temperature, the sum of the different species was normalized to the total concentration (10 μM), using the following equation:

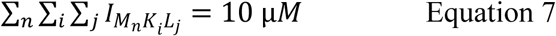
where *I* is the integral of the corresponding peaks obtained from the MS experiment, *M*, *K*^+^ and *L* are the DNA, potassium cations and ligands, respectively, and *n*, i and *j* their stoichiometries. For K^+^ binding to the G-quadruplexes, and for ligand binding to either duplexes or G-quadruplexes, the assumption that response factors are equal is supported by previous studies.^36,38,72^

When a solution containing 100 mM TMAA and 1 mM KCl is sprayed, non-specifically bound K^+^ cations are visible on the peaks of the DNA, whether it is folded or not. To distinguish specific from nonspecific adducts, we used a method described previously.^35,38,73^ Briefly, the distribution of adducts on a reference is used to reconstruct the specific distribution. We checked using the 21 and 22nonG4 sequences (which bind K^+^ only non-specifically) that the distribution of nonspecific adducts is not affected by the temperature (Figure S 29). For each sequence, we used the distribution of adducts obtained at high temperature (80-85 °C) to quantify the contribution of nonspecific adducts to the ion signal, considering that in this temperature range all DNA structures are unfolded. Two nonspecific adducts were taken into account for the 5-charge state and one for the 6-charge state. The detailed mathematical procedure is reported in the supplementary material (Figure S 31).

In Figure S 32, the dotted lines highlight the differences in the melting temperatures obtained for the treatment of each charge state. These differences were negligible (Table S 2). All quantitative results shown in the manuscript are the average of the two charge states. Note that the 4-charge state is also observed but was not included in the summation because the 4-ions present a very high number of nonspecific adducts and would thus bring more uncertainty.^38^

The equilibrium constants are calculated from the ratios of the concentrations, assuming they are equal to the ratios of their activities. The general form of the reactions for which the thermodynamic parameters were extracted (including for duplexes) is given by:

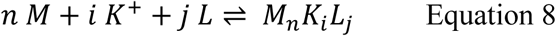

The stoichiometric coefficients *n*, *i* and *j* are not adjustable parameters, but are integer numbers determined unambiguously from the mass spectrometric measurements. The equilibrium constant of the association reactions are thus:

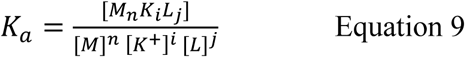

The 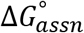 was determined using:

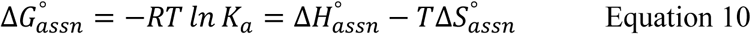
where *R* is the molar gas constant and *T* is the temperature in K. We used the Van’t Hoff plot (ln *K_a_* vs. 1/T) to determine the enthalpic and entropic contributions to the association reaction. The temperature ranges used for the Van’t Hoff plots are reported in the figures of the supplementary material.

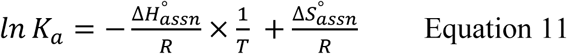

When 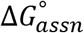 could not be obtained directly at 298K from the measurement, 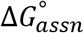 was obtained by extrapolation, using Equation 10 and the uncertainties were calculated from the sum of the uncertainties on 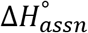 and 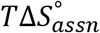.

In most cases the Van’t Hoff plots were linear, and in such cases the 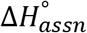 determined from the slope in the temperature range wherein *K_a_* could be determined confidently (nonzero concentration of each component) can be relatively safely extrapolated to other temperatures. In some cases, however, the plots are not perfectly linear, meaning that 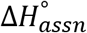 changes with temperature and that 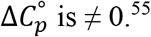. In general, the errors on the linear fits are larger for these cases. Unfortunately, signal flucturations precluded any reliable fitting using quadratic functions.

In cases where the equilibrium constant of a reaction *M* → *M_n_K_i_L_j_* cannot be measured at any temperature because neither species is sufficiently abundant, it is still possible to determine the equilibrium constant and thermodynamic parameters for other coupled equilibria (for example *M* → *M_m_K_k_L_l_* and *M_m_K_k_L_l_* → *M_n_K_i_L_j_*), and then obtain the thermodynamic parameters by difference. The total uncertainty is obtained by standard propagation of individual uncertainties, assuming they are uncorrelated.

## Acknowledgements

The authors thank Marie Paule Teulade-Fichou for providing Phen-DC3 and TrisQ, and Anirban Ghosh for the NMR spectra. The research leading to these results has received funding from the European Research Council under the European Union’s Seventh Framework Programme (FP7/2007-2013) / ERC-2013-CoG 616551-DNAFOLDIMS and FP7-PEOPLE-2012-CIG-333611-BIOPHYMS, from the Inserm [ATIP-Avenir Grant no. R12086GS] and from the Conseil Régional Aquitaine [Grant no. 20121304005].

## SUPPLEMENTARY INFORMATION AVAILABLE

Supplementary MS, CD, NMR spectra and melting data as described in the text is available free of charge via the Internet at http://pubs.acs.org.

The data used in used in this publication will be made freely accessible from the curated data archive of ETH Zurich (https://www.research-collection.ethz.ch) under the DOI: 10.3929/ethz-b-000274776.

## REFERENCES

1. Rhodes, D. & Lipps, H. J. G-quadruplexes and their regulatory roles in biology. Nucleic Acids Res. 43, 8627–8637 (2015).

2. Biffi, G., Tannahill, D., McCafferty, J. & Balasubramanian, S. Quantitative visualization of DNA G-quadruplex structures in human cells. Nat. Chem. 5, 182–6 (2013).

3. Huppert, J. L. & Balasubramanian, S. G-quadruplexes in promoters throughout the human genome. Nucleic Acids Res. 35, 406–413 (2006).

4. Eddy, J. & Maizels, N. Gene function correlates with potential for G4 DNA formation in the human genome. Nucleic Acids Res. 34, 3887–96 (2006).

5. Riou, J.-F. G-quadruplex interacting agents targeting the telomeric G-overhang are more than simple telomerase inhibitors. Curr. Med. Chem. Anticancer. Agents 4, 439–43 (2004).

6. Gomez, D. et al. Telomestatin-induced telomere uncapping is modulated by POT1 through G-overhang extension in HT1080 human tumor cells. J. Biol. Chem. 281, 38721–9 (2006).

7. Rodriguez, R. et al. A novel small molecule that alters shelterin integrity and triggers a DNA-damage response at telomeres. J. Am. Chem. Soc. 130, 15758–9 (2008).

8. Campbell, N., Collie, G. W. & Neidle, S. in Current Protocols in Nucleic Acid Chemistry 17.6.1-17.6.22 (John Wiley & Sons, Inc., 2012). doi:10.1002/0471142700.nc1706s50

9. Webba da Silva, M. NMR methods for studying quadruplex nucleic acids. Methods 43, 264–277 (2007).

10. Adrian, M., Heddi, B. & Phan, A. T. NMR spectroscopy of G-quadruplexes. Methods 57, 11–24 (2012).

11. Li, J., Correia, J. J., Wang, L., Trent, J. O. & Chaires, J. B. Not so crystal clear: the structure of the human telomere G-quadruplex in solution differs from that present in a crystal. Nucleic Acids Res. 33, 4649–59 (2005).

12. Wang, Y. & Patel, D. J. Solution structure of the human telomeric repeat d[AG3(T2AG3)3] G-tetraplex. Structure 1, 263–82 (1993).

13. Parkinson, G. N., Lee, M. P. H. & Neidle, S. Crystal structure of parallel quadruplexes from human telomeric DNA. Nature 417, 876–80 (2002).

14. Luu, K. N., Phan, A. T., Kuryavyi, V., Lacroix, L. & Patel, D. J. Structure of the human telomere in K+ solution: an intramolecular (3 + 1) G-quadruplex scaffold. J. Am. Chem. Soc. 128, 9963–70 (2006).

15. Phan, A. T., Kuryavyi, V., Luu, K. N. & Patel, D. J. Structure of two intramolecular G-quadruplexes formed by natural human telomere sequences in K+ solution. Nucleic Acids Res. 35, 6517–25 (2007).

16. Dai, J., Carver, M., Punchihewa, C., Jones, R. A. & Yang, D. Structure of the Hybrid-2 type intramolecular human telomeric G-quadruplex in K+ solution: insights into structure polymorphism of the human telomeric sequence. Nucleic Acids Res. 35, 4927–4940 (2007).

17. Lim, K. W. et al. Structure of the human telomere in K+ solution: a stable basket-type G-quadruplex with only two G-tetrad layers. J. Am. Chem. Soc. 131, 4301–9 (2009).

18. Wirmer-Bartoschek, J. et al. Solution NMR Structure of a Ligand/Hybrid-2-G-Quadruplex Complex Reveals Rearrangements that Affect Ligand Binding. Angew. Chemie Int. Ed. 56, 7102–7106 (2017).

19. Chung, W. J. et al. Solution structure of an intramolecular (3 + 1) human telomeric G-quadruplex bound to a telomestatin derivative. J. Am. Chem. Soc. 135, 13495–501 (2013).

20. Redman, J. E. Surface plasmon resonance for probing quadruplex folding and interactions with proteins and small molecules. Methods 43, 302–312 (2007).

21. Pagano, B., Mattia, C. A. & Giancola, C. Applications of Isothermal Titration Calorimetry in Biophysical Studies of G-quadruplexes. Int. J. Mol. Sci. 10, 2935–2957 (2009).

22. Rachwal, P. a. & Fox, K. R. Quadruplex melting. Methods 43, 291–301 (2007).

23. Mergny, J. L., Phan, A. T. & Lacroix, L. Following G-quartet formation by UV-spectroscopy. FEBS Lett. 435, 74–78 (1998).

24. Mergny, J.-L. & Lacroix, L. in Current Protocols in Nucleic Acid Chemistry 17.1.1-17.1.15 (John Wiley & Sons, Inc., 2009). doi:10.1002/0471142700.nc1701s37

25. Karsisiotis, A. I. et al. Topological characterization of nucleic acid G-quadruplexes by UV absorption and circular dichroism. Angew. Chem. Int. Ed. Engl. 50, 10645–8 (2011).

26. Pagano, B. et al. Differential scanning calorimetry to investigate G-quadruplexes structural stability. Methods 64, 43–51 (2013).

27. Lim, K. W. et al. Sequence variant (CTAGGG)n in the human telomere favors a G-quadruplex structure containing a G · C · G · C tetrad. Nucleic Acids Res. 37, 6239–6248 (2009).

28. Ambrus, A., Chen, D., Dai, J., Jones, R. A. & Yang, D. Solution structure of the biologically relevant G-quadruplex element in the human c-MYC promoter. Implications for G-quadruplex stabilization. Biochemistry 44, 2048–58 (2005).

29. Do, N. Q. & Phan, A. T. Monomer-Dimer Equilibrium for the 5′-5′ Stacking of Propeller-Type Parallel-Stranded G-Quadruplexes: NMR Structural Study. Chem. - A Eur. J. 18, 14752–14759 (2012).

30. Mergny, J. & Lacroix, L. Analysis of thermal melting curves. Oligonucleotides 537, 515–537 (2003).

31. Gray, R. D. & Chaires, J. B. in Current Protocols in Nucleic Acid Chemistry 17.4.1-17.4.16 (John Wiley & Sons, Inc., 2011). doi:10.1002/0471142700.nc1704s45

32. Gray, R. D., Buscaglia, R. & Chaires, J. B. Populated intermediates in the thermal unfolding of the human telomeric quadruplex. J. Am. Chem. Soc. 134, 16834–44 (2012).

33. Buscaglia, R., Gray, R. D. & Chaires, J. B. Thermodynamic characterization of human telomere quadruplex unfolding. Biopolymers 99, 1006–18 (2013).

34. Bončina, M., Lah, J., Prislan, I. & Vesnaver, G. Energetic basis of human telomeric DNA folding into G-quadruplex structures. J. Am. Chem. Soc. 134, 9657–9663 (2012).

35. Marchand, A. et al. Ligand-induced conformational changes with cation ejection upon binding to human telomeric DNA G-quadruplexes. J. Am. Chem. Soc. 137, 750–756 (2015).

36. Marchand, A., Strzelecka, D. & Gabelica, V. Selective and Cooperative Ligand Binding to Antiparallel Human Telomeric DNA G-Quadruplexes. Chem. - A Eur. J. 22, 9551–9555 (2016).

37. Gabelica, V., Baker, E. S., Teulade-Fichou, M.-P., De Pauw, E. & Bowers, M. T. Stabilization and structure of telomeric and c-myc region intramolecular G-quadruplexes: the role of central cations and small planar ligands. J. Am. Chem. Soc. 129, 895–904 (2007).

38. Marchand, A. & Gabelica, V. Folding and misfolding pathways of G-quadruplex DNA. Nucleic Acids Res. 44, 10999–11012 (2016).

39. Rosu, F., De Pauw, E. & Gabelica, V. Electrospray mass spectrometry to study drug-nucleic acids interactions. Biochimie 90, 1074–1087 (2008).

40. Benesch, J. L. P., Sobott, F. & Robinson, C. V. Thermal Dissociation of Multimeric Protein Complexes by Using Nanoelectrospray Mass Spectrometry. Anal. Chem. 75, 2208–2214 (2003).

41. Wang, G., Abzalimov, R. R. & Kaltashov, I. A. Direct Monitoring of Heat-Stressed Biopolymers with Temperature-Controlled Electrospray Ionization Mass Spectrometry. Anal. Chem. 83, 2870–2876 (2011).

42. El-Baba, T. J. et al. Melting Proteins: Evidence for Multiple Stable Structures upon Thermal Denaturation of Native Ubiquitin from Ion Mobility Spectrometry-Mass Spectrometry Measurements. J. Am. Chem. Soc. 139, 6306–6309 (2017).

43. Hommersom, B., Porta, T. & Heeren, R. M. A. Ion mobility spectrometry reveals intermediate states in temperature-resolved DNA unfolding. Int. J. Mass Spectrom. 419, 52–55 (2017).

44. Alberti, P., Bourdoncle, A., Saccà, B., Lacroix, L. & Mergny, J.-L. DNA nanomachines and nanostructures involving quadruplexes. Org. Biomol. Chem. 4, 3383 (2006).

45. Olsen, C. M., Gmeiner, W. H. & Marky, L. A. Unfolding of G-Quadruplexes: Energetic, and Ion and Water Contributions of G-Quartet Stacking. J. Phys. Chem. B 110, 6962–6969 (2006).

46. Largy, E., Mergny, J.-L. & Gabelica, V. in The Alkali Metal Ions: Their Role for Life (eds. Sigel, A., Sigel, H. & Sigel, R. K. O.) 203–258 (Springer, 2016). doi:10.1007/978-3-319-21756-7_7

47. Miyoshi, D., Karimata, H. & Sugimoto, N. Hydration regulates thermodynamics of G-quadruplex formation under molecular crowding conditions. J. Am. Chem. Soc. 128, 7957–63 (2006).

48. Nakano, S., Miyoshi, D. & Sugimoto, N. Effects of Molecular Crowding on the Structures, Interactions, and Functions of Nucleic Acids. Chem. Rev. 114, 2733–2758 (2014).

49. Bertrand, H. et al. Recognition of G-quadruplex DNA by triangular star-shaped compounds: with or without side chains? Chemistry 17, 4529–39 (2011).

50. Largy, E. et al. Tridentate N-donor palladium(II) complexes as efficient coordinating quadruplex DNA binders. Chemistry 17, 13274–83 (2011).

51. Bertrand, H. et al. Exclusive platination of loop adenines in the human telomeric G-quadruplex. Org. Biomol. Chem. 7, 2864 (2009).

52. Quintana, J. R., Lipanov, A. A. & Dickerson, R. E. Low-temperature crystallographic analyses of the binding of Hoechst 33258 to the double-helical DNA dodecamer C-G-C-G-A-A-T-T-C-G-C-G. Biochemistry 30, 10294–10306 (1991).

53. Graves, D. E. & Wadkins, R. M. in Molecular Basis of Specificity in Nucleic Acid-Drug Interactions (eds. Pullman, B. & Jortner, J.) 177–189 (Kluwer Academic Publishers, 1990). doi:10.1007/978-94-011-3728-7_13

54. Haq, I., Ladbury, J. E., Chowdhry, B. Z., Jenkins, T. C. & Chaires, J. B. Specific binding of hoechst 33258 to the d(CGCAAATTTGCG)2 duplex: calorimetric and spectroscopic studies. J. Mol. Biol. 271, 244–257 (1997).

55. Majhi, P. R., Qi, J., Tang, C.-F. & Shafer, R. H. Heat capacity changes associated with guanine quadruplex formation: An isothermal titration calorimetry study. Biopolymers 89, 302–309 (2008).

56. Chaires, J. B. Possible origin of differences between van’t Hoff and calorimetric enthalpy estimates. Biophys. Chem. 64, 15–23 (1997).

57. Gray, R. D., Trent, J. O. & Chaires, J. B. Folding and unfolding pathways of the human telomeric G-quadruplex. J. Mol. Biol. 426, 1629–50 (2014).

58. Mashimo, T., Yagi, H., Sannohe, Y., Rajendran, A. & Sugiyama, H. Folding pathways of human telomeric type-1 and type-2 G-Quadruplex structures. J. Am. Chem. Soc. 132, 14910–14918 (2010).

59. Monchaud, D. & Teulade-Fichou, M.-P. A hitchhiker’s guide to G-quadruplex ligands. Org. Biomol. Chem. 6, 627–36 (2008).

60. Neidle, S. Quadruplex Nucleic Acids as Novel Therapeutic Targets. J. Med. Chem. 59, 5987–6011 (2016).

61. Bončina, M. et al. Dominant Driving Forces in Human Telomere Quadruplex Binding-Induced Structural Alterations. Biophys. J. 108, 2903–2911 (2015).

62. Bončina, M. et al. Thermodynamic fingerprints of ligand binding to human telomeric G-quadruplexes. Nucleic Acids Res. gkv1167M (2015). doi:10.1093/nar/gkv1167

63. Grand, C. L. et al. The cationic porphyrin TMPyP4 down-regulates c-MYC and human telomerase reverse transcriptase expression and inhibits tumor growth in vivo. Mol. Cancer Ther. 1, 565–73 (2002).

64. Martino, L., Pagano, B., Fotticchia, I., Neidle, S. & Giancola, C. Shedding Light on the Interaction between TMPyP4 and Human Telomeric Quadruplexes. J. Phys. Chem. B 113, 14779–14786 (2009).

65. Ofer, N., Weisman-Shomer, P., Shklover, J. & Fry, M. The quadruplex r(CGG)n destabilizing cationic porphyrin TMPyP4 cooperates with hnRNPs to increase the translation efficiency of fragile X premutation mRNA. Nucleic Acids Res. 37, 2712–2722 (2009).

66. Zamiri, B., Reddy, K., Macgregor, R. B. & Pearson, C. E. TMPyP4 Porphyrin Distorts RNA G-quadruplex Structures of the Disease-associated r(GGGGCC) n Repeat of the C9orf72 Gene and Blocks Interaction of RNA-binding Proteins. J. Biol. Chem. 289, 4653–4659 (2014).

67. Morris, M. J., Wingate, K. L., Silwal, J., Leeper, T. C. & Basu, S. The porphyrin TmPyP4 unfolds the extremely stable G-quadruplex in MT3-MMP mRNA and alleviates its repressive effect to enhance translation in eukaryotic cells. Nucleic Acids Res. 40, 4137–45 (2012).

68. Cavaluzzi, M. J. & Borer, P. N. Revised UV extinction coefficients for nucleoside-5’-monophosphates and unpaired DNA and RNA. Nucleic Acids Res. 32, e13 (2004).

69. De Cian, A., Delemos, E., Mergny, J.-L., Teulade-Fichou, M.-P. & Monchaud, D. Highly efficient G-quadruplex recognition by bisquinolinium compounds. J. Am. Chem. Soc. 129, 1856–7 (2007).

70. D’Atri, V., Porrini, M., Rosu, F. & Gabelica, V. Linking molecular models with ion mobility experiments. Illustration with a rigid nucleic acid structure. J. Mass Spectrom. 50, 711–726 (2015).

71. Darby, R. a J. et al. High throughput measurement of duplex, triplex and quadruplex melting curves using molecular beacons and a LightCycler. Nucleic Acids Res. 30, e39 (2002).

72. Gabelica, V., Rosu, F. & De Pauw, E. A simple method to determine electrospray response factors of noncovalent complexes. Anal. Chem. 81, 6708–15 (2009).

73. Sun, J., Kitova, E. N., Wang, W. & Klassen, J. S. Method for distinguishing specific from nonspecific protein-ligand complexes in nanoelectrospray ionization mass spectrometry. Anal. Chem. 78, 3010–8 (2006).

